# Unreal? A Behavioral, Physiological, and Computational Model of the Sense of Reality

**DOI:** 10.1101/2025.04.07.647542

**Authors:** Gadi Drori, Paz Bar-Tal, Oded Hirsh, Ariel Berlinger, Niva Goldberg, Yair Zvilichovsky, Uri Hertz, Adam Zaidel, Roy Salomon

## Abstract

Our awareness of dreams, hallucinations, and illusions reflects an intriguing human capacity to recognize potentially false perceptions, known as the Sense of Reality (SoR). Though central to mental health, the study of SoR has been hindered by the subjective nature of hallucinatory experiences. Here we employed a novel virtual reality paradigm simulating ‘virtual hallucinations’, mirroring the phenomenology of clinical hallucinations. Combining psychophysics, physiological recordings, and computational modeling in one exploratory (*n* = 31) and one preregistered experiment (*n* = 32), we found that SoR varied with virtual hallucination magnitude and domain. SoR judgments were associated with distinct motor, pupillary, and cardiac responses, allowing classification of virtual hallucination exposure. We present a computational model in which SoR judgments arise from comparing current sensory input to an internal model of the world. Our results shed light on the age-old question ‘how do we know what is real’?

## Introduction

Our ability to function and survive in the world depends on a close correspondence between our representation of the environment and its actual characteristics e.g., (1,2). This correspondence is often termed ‘reality’, and can be contrasted with other states such as dreams and hallucinations which often feel real despite being non-veridical. Thus, while we are typically inclined to trust our sensory experiences, we are also aware that they may not be a veracious representation of the world (3). As such humans continuously monitor the veridicality of their perceptions in a process termed the Sense of Reality (SoR).

The SoR is a central criterion in the assessment of neurological and psychiatric health (4). Distortions of perceptual reality in the form of hallucinations originating from neurological, psychiatric and pharmacological origins are commonplace (5). For example, around 40% of Parkinson’s disease, 27% of schizophrenia patients and between 10-30% of the general population may experience visual hallucinations (6).

Intriguingly, hallucinations can be experienced with or without insight into their nature as false perceptions. For example, patients with visual hallucinations due to macular degeneration (i.e., Charles Bonnet syndrome) typically identify these perceptions as hallucinations (7). Contrarily, psychotic patients often lose the ability to discern between real and hallucinatory percepts, which has been linked to poorer prognosis (8) and associated with reduced cognitive abilities (9). Importantly, lapses in SoR in psychosis have been suggested to arise from a convergence of abnormal perceptual processing and reduced metacognitive monitoring of the reliability of these signals (10). Abnormal SoR may also be relevant to other psychiatric symptoms such as derealization and depersonalization, in which the external world or the self are experienced as strange or unreal (11), typically in the absence of hallucinatory experiences. Despite the obvious clinical significance of SoR, there is little knowledge of the mechanisms supporting its function.

In recent years there is growing evidence suggesting the brain infers the probable causes of incoming sensory information based on learned statistical regularities of the environment (12). Such Bayesian frameworks posit that the brain generates our belief about the source of information (posterior) based on how well it describes the incoming sensory information (likelihood) - and our prior expectations (prior) (13). When a highly atypical sensory event arises (e.g., a rippling building; Fig. 1A) we must evaluate if the cause of this experience is “real” (i.e., a rippling building) or is “unreal” (e.g., I’m hallucinating). Importantly, we constantly encounter slightly atypical perceptual information due to variance in environmental (e.g., unusual reflections or heat haze on a road) and internal (e.g., sensory or neural noise) states. Thus, to infer the source of sensory information we use our prior knowledge of the world (e.g., buildings do not ripple) to decide the probability that this is a veridical perception. Hence, SoR may be considered a process of active inference based on sensory signals and our internal model of the world.

**Figure 1.**
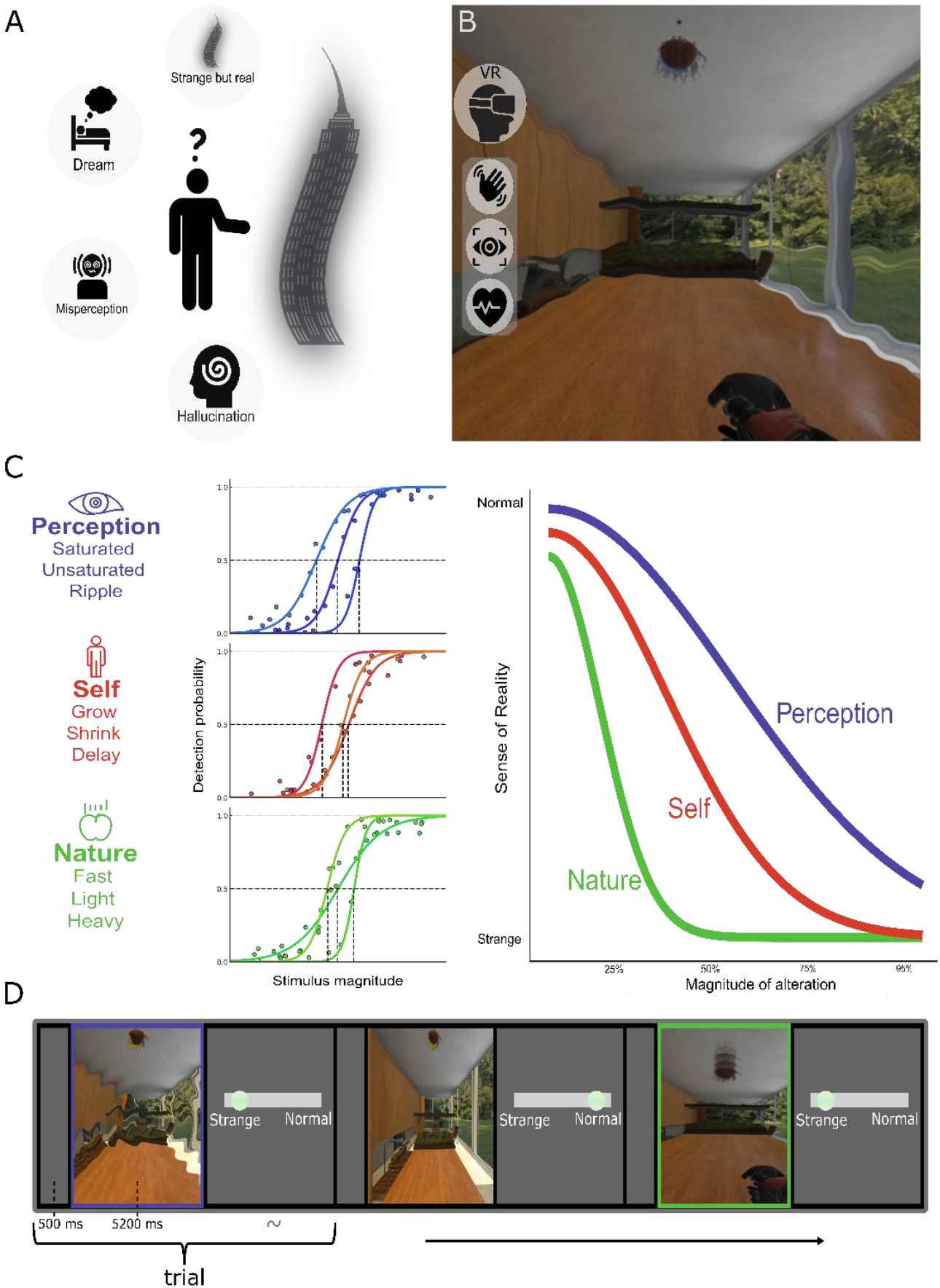
Experimental design. **(A)** Conceptual example of a strange percept requiring a judgment of reality. **(B)** The virtual hallucination paradigm, participants experienced immersive VR scenes with parametrically controlled distortions while physiological responses (kinematics, pupil size, ECG) were recorded. **(C) Left**: virtual hallucinations targeted three domains with distinct real-world likelihoods (Perception, Self, and Nature). **Middle**: Psychophysical modeling matched salience across domains and participants using four graded magnitudes based on detection thresholds (baseline = 0%, 1^st^ = 25%, 2^nd^ = 50% (the JND), 3^rd^ = 75%, 4^th^ = 95%). **Right**: These detection thresholds were used as the stimulus magnitudes in 360 randomly ordered SoR rating trials across domains. To control for potential drifts in ratings as a result of the participants losing touch with the baseline condition, we placed 5 baseline trials every 40 virtual hallucination trials. **(D)** Three example SoR rating trials, with colored frames indicating trial type (blue = Perception, green = Nature, black = baseline). Participants rated each trial from “normal” to “strange” (100–0), quantifying subjective deviations in reality. Each trial lasted 5.2 seconds, during which a dynamic event prompted a reaching response from the participant. This design enabled controlled assessment of domain and magnitude-specific SoR effects across behavioral and physiological signals.

Such causal inference approaches have been successful in modelling many aspects of human perception and behavior (13). Under this framework, SoR can be viewed as a probabilistic inference expressing the fit between a given sensory signal and our model of the world. For example, sudden changes in the color spectra of the world could be attributed to a wide range of causes (e.g., lighting changes, colored glasses) which we encounter regularly (i.e., wide likelihood distribution). As a result, changing colors of the visual scene, for which we have more experience, should have lower impact on SoR. Contrarily, alterations of aspects of reality for which we have previously experienced little variation should have higher impact on our SoR. For example, changing sensorimotor contingencies (e.g., a delay between actions and their consequences) or the laws of nature (e.g., changes in gravity) for which we normally have little experience (i.e. narrow likelihood distribution) (14). Taking this perspective, we suggest that SoR inferences are grounded in the learned regularities of the environment.

Measuring ecologically valid reality judgements in response to non-veridical perceptions such as hallucinations presents significant methodological challenges. Typically confined to psychedelic, psychiatric or neurological states (15,16), hallucinations are notoriously difficult to study scientifically due to their transient, ineffable and subjective nature (17,18). Previous related work has therefore focused on analogous phenomena. For example, reality monitoring studies (also known as source monitoring), have investigated the ability to distinguish internally from externally generated information (i.e., whether remembered information originated from perception or imagery) by evaluating characteristics such as vividness or sensory detail (19–21). In another line of work, a recent study utilized the motion-aftereffect illusion to probe perceptual insight by dissociating perceptions from beliefs about stimuli (22). While informative, these approaches do not capture key aspects of the visual phenomenology of hallucinations, particularly the feeling of strangeness or “bizarreness” that is central to understanding aberrant SoR processing. Accordingly, studying SoR requires a replicable, quantitative approach capable of simulating such experiences in controlled settings.

To address these challenges, we developed an immersive Virtual Reality (VR) paradigm designed to simulate visual hallucinations under controlled experimental conditions. We selected several aspects of reality to be examined in this project, based on the phenomenology of distorted visual reality characteristic of hallucinatory states (15,23). These ‘virtual hallucinations’ broadly fall into three domains: (1) Changes in ‘Perception’, in which the visual appearance of the scene is manipulated (e.g., colors, acuity). Such perceptual changes are well documented in hallucinations arising from pharmacological or medical origins (16) and are a central feature of schizotypy (24). (2) Laws of ‘Nature’, in which visual representations of physical laws (e.g., gravity, passage of time) are altered. Changes in the experience of natural physics are common during dreaming (25), and are a defining feature of psychedelic and mystical experiences (26). (3) Changes of ‘Self’, in which the participants’ sense of self is disturbed through conflicts between visual signals and self-related information (e.g., sensorimotor conflicts, changes in first person perspective) (27). Changes in the experience of the self are a hallmark of altered reality in neurological (28), psychiatric and mystical contexts (29). Importantly, the VR environment allows precise manipulation of these domains, enabling us to alter reality at qualitatively different scales (Fig. 1B-D). For example, we can change the participants’ first-person perspective from floor level to ceiling height, simulating dramatic changes of self that occur in hallucinatory states (e.g., Alice in Wonderland Syndrome (30)). We can also introduce subtle changes, barely noticeable to participants, resembling prodromal or derealization-like experiences (11,31).

Critically, we used psychophysics to individually calibrate the intensity of virtual hallucinations, enabling comparison of subjective experiences across different magnitudes and conditions, within and between participants. We present evidence from two experiments targeting SoR, an initial exploratory experiment - Experiment 1, followed by a pre-registered replication - Experiment 2 (see pre-registration: https://osf.io/bt3kd). Our behavioral results are accompanied by converging evidence from physiological measures (kinematics, pupillometry and cardiac activity) demonstrating physiological responses to manipulations of SoR. To further characterize the mechanisms of SoR, we utilized a Bayesian inference model providing new insights into the perceptual decision process underlying reality judgments.

### Pre-registered hypotheses

We pre-registered our hypotheses, study design, sampling plan, and analysis methods based on findings from Experiment 1 prior to analyzing the data from Experiment 2.

Hypothesis 1 (H1). Subjective judgments:

a. Subjective SoR (mean of ratings) will diminish with increasing virtual hallucination magnitude.
b. Changes in SoR ratings between virtual hallucination magnitudes will vary depending on domain.

Hypothesis 2 (H2). Physiological signals:

a. Hand kinematics during the reaching phase will vary between *altered* and *unaltered* trials.
b. Hand kinematics during the reaching phase will vary between *altered* and *unaltered* trials.
c. Pupillometric data will vary between *altered* and *unaltered* trials.
d. Cardiac activity will vary between trials subjectively experienced as *strange* or *normal*.
e. Cardiac activity will vary between domains in *strong* trials (3^rd^ & 4^th^ magnitudes).

Hypothesis 3 (H3). Clinical correlations:

a. We hypothesized that there will be correlations between the slope of ratings in the Self domain and total scores of self-reported clinical measures of abnormal-self and abnormal-perception (see Supplementary Materials “Clinical measures”).

### Notation guide

The relationships of reported results to pre-registered hypotheses are indicated as follows - † Indicates a fully confirmed hypothesis; ‡ indicates a partially confirmed hypothesis; × indicates an unconfirmed hypothesis. Where applicable, the specific hypothesis number is referenced (e.g., [H1.a †]).

## Results

The analyses reported below were pre-registered, unless otherwise specified. Unregistered exploratory analyses are introduced separately toward the end of the results section (see “Classifying SoR from physiological signals” and “SoR ratings fit to a Bayesian causal inference model”).

### SoR varies across domains and magnitudes

Analysis using mixed-effect models to predict SoR ratings revealed, as hypothesized, a main effect for virtual hallucination magnitude in Experiment 1 (*F*4, 11,012 = 683.5, *p* < 0.0001, η^2^ = 0.2) and Experiment 2 (*F*_4, 11,391_ = 537.7, *p* < 0.0001, η^2^ = 0.16), confirming that SoR ratings decreased with increasing magnitude of alteration [H1.a †].

There was a main effect of domain in Experiment 1 (*F*_2, 30_ = 12.3, *p* < 0.0001, η^2^ = 0.44; Fig. 2), reflecting a tendency to rate Nature (*z* = −4.5, *p* < 0.0001, *d =* −0.3) and Self (*z* = 2.7, *p* = 0.016, *d =* 0.14) virtual hallucinations as less real than Perception (Nature: 61.4 ± 34.2, Self: 67.2 ± 31.5, Perception: 72.6 ± 30.1). This effect was not significant in Experiment 2 (*F*_2, 31_ = 2.2, *p* = 0.12; Nature: 61.1 ± 34.2, Self: 63.1 ± 34.4, Perception: 64.7 ± 33.7). However, as predicted, an interaction was found between magnitude and domain both in Experiment 1 (*F*_8, 11,012_ =20.1, *p* < 0.0001, η^2^ = 0.01) and Experiment 2 (*F*_8, 11,391_ = 6.2, *p* < 0.0001, η^2^ = 0.06) [H1.b †]. Together these results show that the changes in SoR ratings across magnitudes depended on virtual hallucination domain (Fig. 2A and Fig. S1).

**Figure 2.**
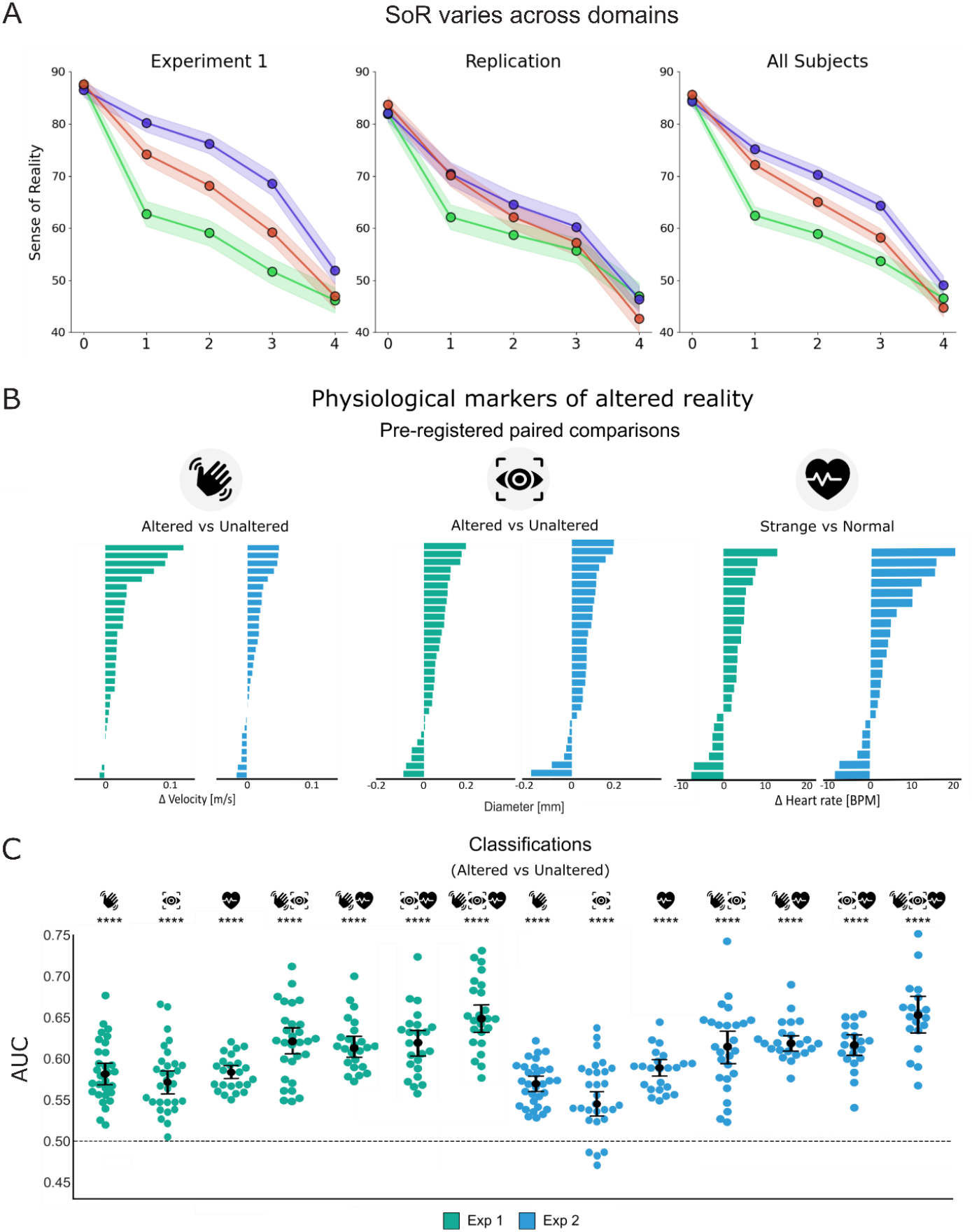
SoR varies across domains. **(A)** SoR ratings by virtual hallucination magnitude and domain for Experiment 1 (*n* = 31), Experiment 2 (*n* = 32), and combined data (*N* = 63). Lines represent group averages for Perception (blue), Self (red), and Nature (green), the colored dots represent group means per magnitude, and the shaded ribbons represent the 95% CI. A mixed-effects model revealed a significant effect for magnitude and a domain × magnitude interaction in both experiments. **Physiological markers of altered reality. (B)** Physiological responses for each participant, based on paired comparisons from the pre-registered analysis plan. Each bar represents the relative change per subject in kinematic (*n*_1_ = 30, *n*_2_ = 30) **(Left)**, pupillometric (*n*_1_ = 30, *n*_2_ = 29) **(Middle)**, and cardiac (*n*_1_ = 24, *n*_2_ = 23) **(Right)** measures between trial types. **Classifying SoR from physiological signals. (C)** Single-subject classification performance (AUC) for predicting *altered* vs. *unaltered* trials using kinematic, pupillometric, and cardiac measures. Each dot represents an individual participant; error bars show standard error, and the dashed line indicates chance level (AUC = 0.50). Classification for each unique signal was significantly above chance in both experiments (Exp 1: AUC = 0.58; Exp 2: AUC = 0.55–0.59, all *p* < 0.0001). Dual-modality combinations improved classification accuracy, reaching AUC ≈ 0.62 (Exp 1) and 0.60– 0.61 (Exp 2), while three-signal combinations achieved the highest performance (AUC = 0.65 and 0.64 for Exp 1 and Exp 2, respectively).

To examine whether individual differences in SoR ratings related to clinical symptoms, we assessed anomalous perceptual experiences using the CAPS, PQ-B, and IPASE questionnaires. Participants reported low, non-clinical scores overall, and contrary to our hypothesis, SoR ratings did not correlate with questionnaire scores in any domain (all Spearman’s ρ ≤ 0.20, all *p* ≥ 0.13) [H3.a ×].

### Physiological markers of altered reality

To understand physiological responses to altered reality, we analyzed hand kinematics, pupil dilation, and heart rate, comparing them by virtual hallucination presence (i.e., *altered* vs. *unaltered* trials) and subjective experience (i.e., *strange* vs. *normal* trials) (see Materials and Methods “Statistical analysis”). Exposure to virtual hallucinations elicited robust physiological responses across both experiments.

Analysis of kinematic data from Experiment 1 revealed a difference in movements during virtual hallucinations. Specifically, mean hand velocity during the reaching phase was higher in *altered* compared to *unaltered* trials (0.47 ± 0.16 m/s vs. 0.44 ± 0.17 m/s; 13.48% increase; *W*_29_ = 23, *p* < 0.0001, *r* = 0.71) and in trials subjectively experienced as *strange* compared to trials experienced as *normal* (0.48 ± 0.16 m/s vs. 0.45 ± 0.17 m/s; 17.08% increase; *W*_29_ = 33, *p* < 0.0001, *r* = 0.71). These effects were replicated in Experiment 2 (*altered* vs. *unaltered*: 0.39 ± 0.14 m/s vs. 0.37 ± 0.15 m/s; 8.85% increase; *t*_29_ = −3.59, *p* < 0.0001, *d* = 0.65) [H2.a †]; (*strange* vs. *normal*: 0.39 ± 0.14 m/s vs. 0.37 ± 0.15 m/s; 13.82% increase; *t*_29_ = −5.11, *p* < 0.0001, *d* = 0.93) [H2.b †]. Thus, objective and subjective aspects of virtual hallucinations were both associated with significant changes in sensorimotor behavior (Fig. 2B & S2).

Pupil size changed significantly during virtual hallucinations. The effect was evident in the 4–5 s window of Experiment 1 (Δ 0.57 ± 0.18 mm; *t*_29_ = 4.15, *p* < 0.0001, *d* = 0.76) and was replicated in Experiment 2 (Δ 0.67 ± 0.22 mm; *t*_28_ = 3.78, *p* < 0.0001, *d* = 0.7) [H2.c †]. Additional analysis revealed the effect remained significant when comparing over the entire trial duration (0-5 s) in both experiments: Experiment 1 (Δ 0.36 ± 0.12 mm; *t*_29_ = 3.18, *p* < 0.001, *d* = 0.58); Experiment 2 (Δ 0.41 ± 0.16 mm; *t*_28_ = 2.55, *p* < 0.001, *d* = 0.47), demonstrating a robust association between pupil dilation and virtual hallucination presence (Fig. 2B).

Heart rate was found to decelerate during trials experienced as *strange* compared to trials experienced as *normal* in Experiment 1 (χ^2^_1_ = 4.38, *p* = 0.036, *R*^2^_c_ = 0.39, *R*^2^_m_ = 0.09) and Experiment 2 (χ^2^_1_ = 15.3, *p* < 0.0001, *R*^2^_c_ = 0.79, *R*^2^_m_ = 0.09) [H2.d †]. Interestingly, in *strong* trials heart rate deceleration showed a significant domain effect in Experiment 1 (χ^2^_2_ = 14.63, *p* < 0.001, *R*^2^_c_ = 0.37, *R*^2^_m_ = 0.055), which was most pronounced in Nature (−0.38 ± 0.1), followed by Self (−0.21 ± 0.1), and Perception (Fig. S3). However, this effect did not replicate in Experiment 2 (χ^2^_2_ = 5.82 *p* = 0.054) [H2.e ×]. Leading us to conclude that heart rate deceleration was only consistently associated with the subjective strangeness of the trial, but not with virtual hallucination domain.

### Classifying SoR from physiological signals

The following analyses were conducted as an exploratory extension beyond our pre-registered analyses to assess whether physiological measures could predict SoR ratings (see Materials and Methods “Classification of physiological signals”). Machine learning classifiers revealed significant, above-chance trial prediction using individual signals (0.55 ≤ AUC ≤ 0.59), with higher accuracy for combinations of signals (0.60 ≤ AUC ≤ 0.65) (Fig. 2C). Notably, a cross-subject model trained on pupil data from Experiment 1 successfully classified virtual hallucination presence in Experiment 2 (AUC = 0.53 ± 0.03, *t*_28_ = 4.99, *p* < .001, *d* = 1) (Fig. S4A). These results indicate that physiological signals contain objective and subjective information about SoR, with each signal contributing unique variance and their combinations improving predictive accuracy.

### SoR ratings fit to a Bayesian causal inference model

The computational model analysis was conducted post hoc on the combined data of both experiments to quantify the inference processes underlying SoR ratings. Our model assumes the observer infers the category of an unusual event as real or unreal by deriving the log ratio of the two possibilities given the evidence and their prior expectations. The model predictions effectively simulated the observed domain differences (Figs. 3A-B). We examined the domain effect in the model’s predicted ratings by analyzing three parameters underlying reality judgments: *p*_*real*_ the prior probability that an event is real; and σ, the expected sensory noise level. Together these values determine a third parameter the decision criterion (*k*), representing an evidence threshold for reality judgments (see Methods for full computational model description).

**Figure 3.**
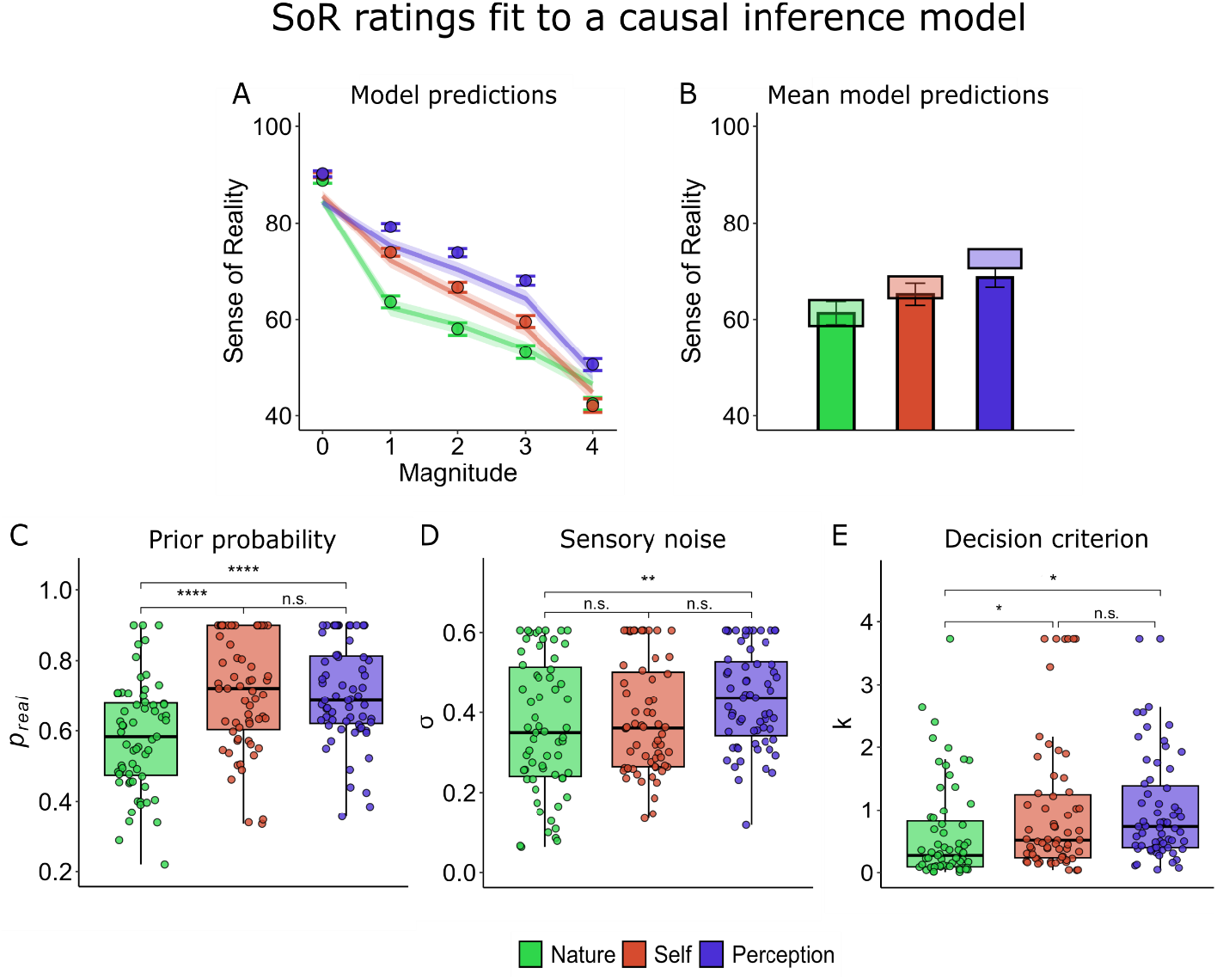
SoR ratings fit to a Bayesian causal inference model. **(A)** Model predictions (points) overlayed on the group-average SoR ratings (*n* = 63). **(B)** Observed (bars represent the mean, error bars represent the 95% CI) versus predicted ratings (tiles, tile size and location represent the mean ± 2 SE). **(C) Right**: *p*_*real*_ was lowest for Nature, indicating lower baseline plausibility; **Middle**: The estimated sensory noise (σ) was lowest for Nature, reflecting more precise expectations; **Left**: Decision criterion (*k*) was strictest for Nature, suggesting greater sensitivity to violations.

Each parameter was analyzed using a separate linear mixed-effects model with domain as a fixed effect and participant as random intercept. The analyses revealed a significant main effect of domain for *p*_*real*_ (*F*_2_ = 17.9, *p* < 0.0001, η^2^ = 0.22); σ (*F*_2_ = 5.3, *p* < 0.01, η^2^ = 0.08) and *k* (χ^2^_2_ = 9.7, *p* < 0.01, *R*^2^_c_ = 0.3, *R*^2^_m_ = 0.04), indicating that domain systematically influenced the likelihood and decision criterion for real events (Fig. 4C). Post-hoc pairwise comparisons using estimated marginal means with Tukey correction revealed that Nature had the lowest *p*_*real*_ (0.58 ± 0.15), significantly different from Perception (0.7 ± 0.14, *t*_124_ = −5, *p* < 0.0001, *d* = −0.63) and Self (0.71 ± 0.16, *t*_124_ = 5.3, *p* < 0.0001, *d* = 0.67), suggesting that participants were biased to assume the least prior probability for Nature virtual hallucinations being real. Similarly, Nature had the lowest σ, with a significant difference observed between Nature and Perception (*t*_124_ = −3.15, *p* < 0.01, *d* = −0.56), reflecting a narrower likelihood distribution for Nature and a wider one for Perception (Nature: 0.36 ± 0.16, Self: 0.38 ± 0.14, Perception: 0.43 ± 0.12). Finally, Nature exhibited the strictest decision criterion *k* (0.49 ± 0.78), significantly more conservative than Self (0.74 ± 1.1; *z* = 2.4, *p* = 0.04, *d* = 0.22) and Perception (0.88 ± 0.85; *z* = −3.8, *p* < 0.001, *d* = −0.35), reinforcing that domain differences in SoR judgments stem from different prior expectations of what could be real.

**Figure 4.**
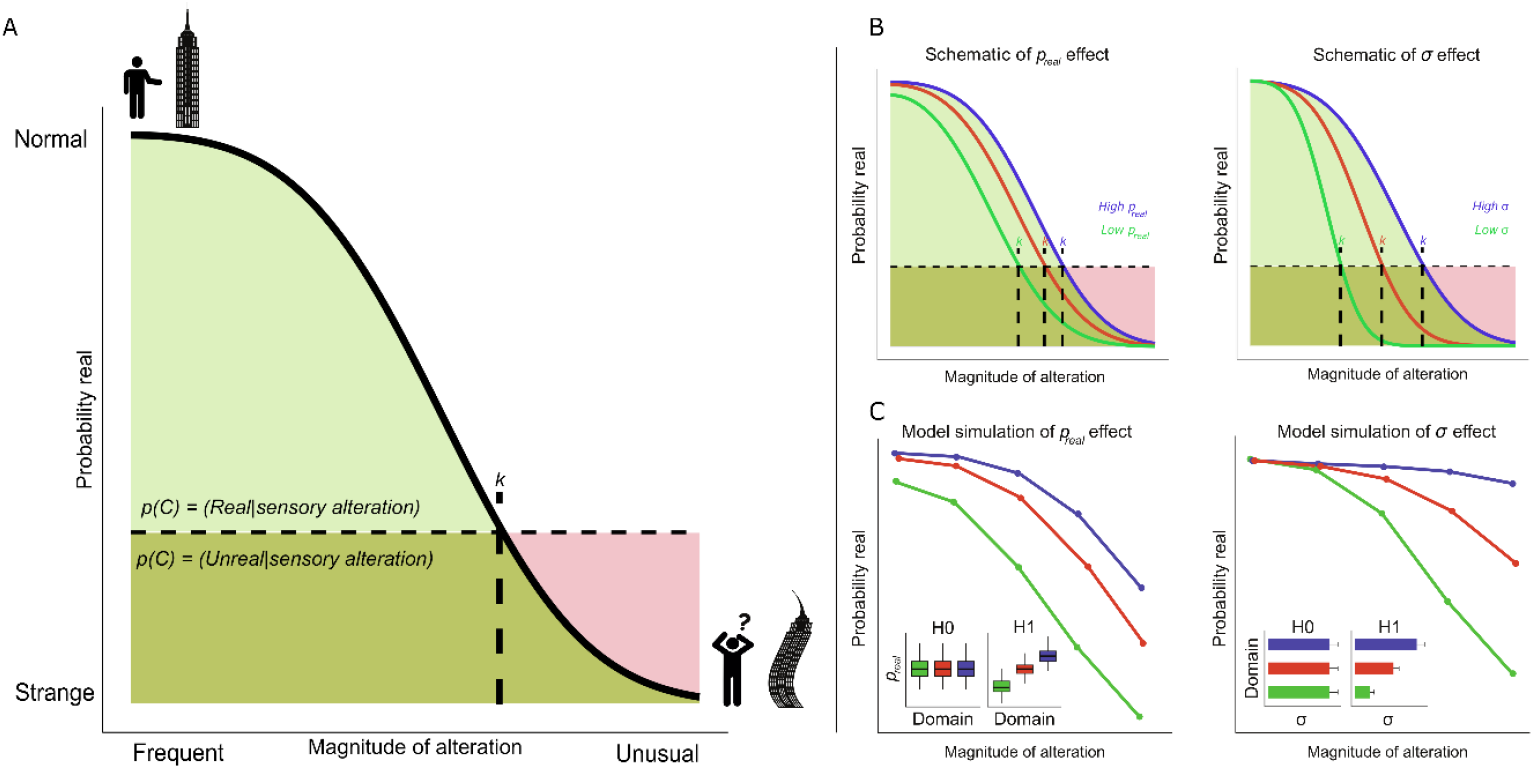
Computational model of SoR. Illustration of the proposed inference model of SoR. The model incorporates prior expectations for the probability and perceptual uncertainty of real and unreal events, leading to dynamic decision thresholds. **(A)** The decision to classify an event as real or unreal is determined by the log-posterior ratio of two competing posteriors: *p*(*C* = Real | *x*_*trial*_) and *p*(*C* = Unreal | *x*_*trial*_). The real posterior is modeled as a half gaussian (light green), the unreal posterior is modeled as a wide, shallow half-gaussian (light red), representing a generalized belief that all experiences could be unreal, especially extreme deviations from the norm. The posteriors overlap where both hypotheses assign probability to the observed event (dark green). The threshold *k* is defined as the point of equal likelihood (i.e., where the green and red distributions intersect). Smaller scale, more frequent events fall to the left of *k* and are more likely judged as real, while larger scale, more unusual events fall to the right and are more likely judged as unreal. **(B)** Illustration of how *p*_*real*_ and σ affect *k*. **Left:** *p*_*real*_ = 0.9 (blue), 0.8 (red), 0.6 (green). Higher *p*_*real*_ leads to more conservative judgments, especially for weaker events. **Right:** σ = 0.7 (blue), 0.6 (red), 0.4 (green). The likelihood of events being judged as real increases with the expectation of sensory noise. **(C)** Model simulation showing how varying *p*_*real*_ and σ shapes predicted SoR judgments.

## Discussion

In this study, we employed an ecological virtual reality paradigm designed to simulate the phenomenology of visual hallucinations and systematically investigate the Sense of Reality (SoR). Our primary findings reveal that: (1) virtual hallucinations from different domains - Nature, Self, and Perception - have distinct and replicable effects on reality judgments, with violations in the laws of nature being perceived as the least real, followed by those related to the self and perception. (2) Experiencing virtual hallucinations was associated with consistent physiological changes in kinematics, pupil size, and cardiac activity. (3) These findings were further supported by a perceptual inference model, which successfully captured the domain-specific effects by accounting for differences in prior expectations for the likelihood of sensory evidence.

### Reality judgments differ across domains

We used a novel approach in which we simulated in virtual reality the reported phenomenology of nine different types of hallucinations to study their effects in controlled conditions. Utilizing psychophysical staircases, we controlled the subjective saliency of these different virtual hallucinations and demonstrated that SoR ratings significantly differ across domains (Perception, Self, Nature), with each domain showing distinct effects on participants’ reality judgments. Notably, alterations in the Nature domain were experienced as the least real, followed by alterations in the Self and Perception domains. This finding aligns with predictive processing theories (32), which suggest that violations of stable environmental regularities (such as those pertaining to the laws of nature) would lead to a larger disruption in SoR. For example, changes in the perceptual domain such as deviations in color saturation (reported in depression (33) and psychedelic (34) hallucinations) effected reality judgments less than deviations in gravity. In light of this, we propose that SoR judgments are based on accumulated prior experience of the observer, thus domains for which a wider range of experiences (wide likelihood distributions) are available will allow more flexibility before inducing a feeling of disturbed reality.

### Physiological correlates of altered SoR

Our study provides robust evidence that exposure to virtual hallucinations not only affects subjective reality judgments but also induces changes in physiological responses. Our observation of consistent modulations in hand kinematics, pupil size and cardiac activity, during virtual hallucinations underscore the interplay between predictive mechanisms of perception and the body’s sensorimotor and autonomic functions.

During virtual hallucinations hand velocity was higher, allowing to classify trial by trial exposure to altered reality states well above chance level. Modulation of sensorimotor processes are well documented in psychiatric and neurological disorders involving hallucinatory experiences such as Parkinson’s disease and schizophrenia (35,36). We suggest that the violated sensory expectations elicited by the virtual hallucinations may have manifested as a more automatic and rapid reaching movement, possibly related to affective sensations of uncanniness (37,38). Consistent with this, hand kinematics enabled classification of not only objective exposure to virtual hallucinations but also subjective feelings of disturbed reality (Fig. S2). Thus, kinematic changes offer a non-invasive, objective marker of altered reality states and their impact on motor function.

Cardiac responses decelerated significantly during trials subjectively experienced by the participants as *strange*, consistent with past findings linking cardiac responses to changes in the sense of self, consciousness (39,40), psychedelic hallucinations (41) and perceptual inference (42,43). Specifically, cardiac deceleration predicted subjective feelings of strangeness and may be a parasympathetic response to mismatches between sensory input and internal predictions (42). Pupil dilation showed a clear and replicable response to virtual hallucinations. Pupillometry, correlated with norepinephrine release (44), serves as a direct and implicit measure of various internal cognitive processes (45), including prediction errors (46). Here, the observed pupil dilations may reflect responses to the cognitive effort required to process unpredicted sensory information. Critically, pupillometry classified altered trials both within and across participants indicating a distinct yet generalizable pupil response to reality alterations (Fig. S4A). Moreover, pupil dilation increased with alteration magnitude, suggesting it gauges prediction violations (Fig. S4B-C). This marker provides a real-time, non-invasive measure of SoR, with potential to inform the detection and clinical assessment of hallucination severity.

The physiological markers identified here offer a valuable opportunity to advance the study and assessment of hallucinations, providing an objective complementary approach for continuous, real-time monitoring of hallucination-related states without relying solely on traditional self-reports, which are inherently subjective. This approach is particularly advantageous in clinical settings where patients may have limited insight into their hallucinations or may be unable or unwilling to communicate their experiences. By providing a quantifiable and reproducible method to detect and track hallucinations, these measures could supplement clinical assessments, enabling more accurate diagnosis and monitoring of conditions such as schizophrenia, Parkinson’s disease, and other disorders involving perceptual disturbances.

### A perceptual inference model of SoR

Our computational model demonstrates how sensory expectations, such as the prior probability or the expected sensory noise assigned to an event, systematically influence the decision rule for reality judgments. Importantly, the main strength of the model lies in its ability to explain the differences in SoR judgments between domains in terms of differently biased sensory expectations. This pattern is consistent with previous findings (47), and reinforces the idea that the Sense of Reality depends on prior expectations shaped by the statistical regularities of the environment.

While the model provides a compelling fit to the data, we do not claim it to be the definitive or most complete model for SoR. Rather, it serves as a preliminary conceptual model, demonstrating how domain differences in SoR judgments can emerge from variations in priors and likelihoods. Future modeling efforts should incorporate additional factors such as learning, attention, and the influence of clinical symptoms (48), which will likely contribute to a more nuanced and dynamic understanding of the SoR.

Despite its conceptual nature, the current framework raises key questions about how perceptual reality priors differ across individuals and conditions. For instance, are such priors different in pathology? Abnormal predictive processing has been repeatedly implicated in psychosis and related disorders, where irregular weighting of sensory evidence or priors may disrupt the capacity to distinguish real from unreal events (49). Extending the present computational model to clinical populations may enrich our understanding of how predictive abnormalities relate to symptoms such as psychotic hallucinations and derealization (50). Such models may also inform new approaches for clinical assessment, by linking deviations in SoR to quantifiable parameters of perceptual inference (51). Early detection of psychotic symptoms has been linked to improved prognosis (52), an equally important question concerns how perceptual reality priors are shaped over the course of development? Children are more prone to psychotic-like experiences, which have been suggested to represent early markers of risk for psychosis (53). SoR is not impaired during childhood, yet seems qualitatively different from adults, likely reflecting a combination of inexperience (54) with cognitive and sensory immaturity (55,56). Longitudinal investigations of SoR across development could inform us on how internal models of reality are shaped and when they become vulnerable. Finally, our experimental approach significantly facilitates the capacity for neuroimaging of hallucination-like states. Further studies using fMRI and EEG could help identify the neural mechanisms underlying SoR.

### Limitations

A potential limitation of this study is that participants were fully aware they were in a virtual environment and may have therefore treated all experiences within it as ‘unreal’ by default. However, this concern is likely outweighed by the highly immersive nature of VR, which enables strong bottom-up simulations that can elicit realistic perceptual and physiological responses, even when participants know the experience is artificial (57,58). In evidence of this, our paradigm produced large changes in subjective reality ratings (η^2^ ≥ 0.16) and moderate to large physiological responses (*d* ≈ 0.5-0.9), suggesting that the virtual hallucinations successfully simulated unusual experiences. The current study used only two sensory modalities to test SoR (visual and sensorimotor), future research should extend SoR paradigms to other relevant scenarios, such as auditory hallucinations, feelings of presence and multimodal hallucinations which are common symptoms of schizophrenia (59,60). Moreover, the absence of correlations with psychosis symptoms in non-clinical samples emphasizes the need for testing SoR in clinical populations.

In conclusion, we developed a novel experimental approach allowing to rigorously test the Sense of Reality by simulating hallucinations. Our results demonstrate how Sense of Reality judgments are dependent on internal models of the world, and how experiences deviating from these models induce changes in phenomenology and physiological markers. This framework significantly advances our understanding of hallucinatory experiences and may advance tools for assessing and treating clinical symptoms of altered SoR.

## Materials and Methods

### Experimental design

We implemented a within-subject design in which all participants followed the same experimental protocol, to assess how SoR is influenced by the domain and magnitude of virtual hallucinations. Each participant attended two sessions. Session 1 was devoted to measuring individual psychophysical thresholds, which were used as normalized condition magnitudes for Session 2. In the second session, participants completed the SoR rating task, consisting of 360 randomly ordered trials varying in domain (Perception, Self, Nature) and magnitude (five levels based on detection thresholds). In both sessions participants sat in front of a table at the center of the experiment room and wore a VR headset and an ECG harness. After the task participants filled out three self-report questionnaires assessing traits related to abnormal perception.

### Participants

63 participants were recruited for two experiments. Experiment 1 included 34 participants, of which 2 dropped out during the experiment and 1 was excluded due to insufficient data, leaving a total of 31 (mean age 27.6 years ± 5.3 years, 14 females). Based on the effect sizes found in Experiment 1, we performed a power analysis that indicated a minimum sample size of 32 was required to replicate the effects (Supplementary Materials “Power analysis”). For Experiment 2, we recruited 40 participants, of which 8 were excluded whose responses were not sufficiently captured by psychometric fits (see pre-registration and Supplementary Materials “Data exclusion criteria”), resulting in a final sample of 32 (mean age 25.1 ± 4.5 years, 22 females). All participants provided written informed consent (IRB ISU202012001), were right-handed with normal or corrected to normal vision, and self-reported no substance dependence and no current or prior history of significant medical or neurological illness or use of psychiatric medication.

### Virtual reality hardware and environment

During the experiments participants wore an HTC Vive-pro Eye head-mounted display (HMD) with eye-tracking and retractable headphones and responded using a touch-sensitive hand controller (61). The baseline virtual environment was constructed as a furnished house interior, providing a normative setting for subsequent reality manipulation using virtual hallucinations. We enhanced the realism of the environment by matching several of the task-relevant virtual items with real ones, thus adding physical correspondence to the experience (Supplementary Materials “Setup and calibration procedures”). Inside the VR environment the participants found themselves positioned at the center of the virtual house in front of a corresponding virtual table. From their seated position they were able to look around and interact with the environment. The matching dimensions of the real and virtual tables amplified the tactile, proprioceptive and visual feedback of the participants exact position and height in VR. Their right hand and controller were represented by a virtual hand and an identical virtual controller. The virtual controller and hand provided similar feedback on the correspondence between real and virtual hand motions. To ensure engagement and attention to the virtual environment, on each trial participants performed a sensorimotor task in which they had to point to a virtual butterfly which could appear at several locations (see “Secondary Task”).

### Secondary Task

In each trial, participants observed a dynamic event during which a hidden butterfly briefly appeared, requiring them to reach toward it with their right hand before returning to a central start position. This task ensured that participants continuously attended to the environment and engaged with its visual appearance, physical dynamics, and first-person perspective. After the scene ended, participants reported their subjective impression of the environment on a sliding scale ranging from “normal” to “strange”. For a complete description see Supplementary Materials “Detailed secondary task”.

### Experimental Conditions

We leveraged the properties of VR to induce specific, highly controlled alterations in the environment, mirroring phenomena reported in psychiatric and neurological conditions (5,28,62). Three theoretical domains of experience were targeted: Nature, Self, and Perception. In the Nature domain, we modified the physics of object motion by adjusting the gravitational constant and temporal rate, producing ‘heavy’ and ‘light’ gravity as well as ‘fast’ time-perception virtual hallucinations, making objects appear to move at the wrong speed, or fall in unexpected ways. In the Self domain, we altered participants’ first-person perspective by shifting camera height (‘grow*’* and ‘shrink’ conditions) and perturbed their sense of embodiment and agency by introducing a lag between real and virtual hand movements (‘delay’ condition). In the Perception domain, we manipulated visual properties using filters that enhanced or reduced color saturation (‘saturated’, ‘unsaturated’), or distorted the texture of the scene (‘ripple’, a peripheral visual warping effect).

## Experimental Procedures

### Psychophysics experiment

To ensure valid comparisons across virtual hallucination conditions and between participants, we calibrated stimulus intensities individually using a two-alternative-forced-choice (2AFC) 1-up\1-down descending staircase procedure (Fig. 1C, Middle). The order of conditions was randomized between participants. Participants viewed two sequential phases on each trial, one containing a virtual hallucination, and reported whether they detected a difference. Stimulus intensity was adjusted based on detection performance until meeting acceptable fit criteria (see Supplementary Materials “Detailed staircase procedure” for full methodology).

### Instructions and training for psychophysics experiment

Participants were instructed on the staircase procedure, VR equipment and environment before entering VR. Participants were then introduced to the VR equipment, and environment to familiarize themselves with the unaltered virtual environment. Once in VR participants were instructed again in the psychophysical staircase task. Participants trained before the task with a structured staircase as an example. On each trial, participants were immersed in the virtual environment while performing the secondary task for two phases (5.2 s each), one of which contained a virtual hallucination. After viewing both phases the participant was required to judge if both phases felt the ‘same’ (press right) or ‘different’ (press left).

### Psychophysical curve estimation

For psychometric-curve estimation we adapted the procedure from Feigin et al., 2021 (63). We fitted the data from the staircase procedures per participant in all conditions to a logistic function using the Matlab Psignifit toolbox (64), retaining only fits with pseudo-*R*^2^ ≥ 0.3 (Supplementary Materials “Detailed curve estimation procedure”). We then extracted four detection thresholds for each virtual hallucination from the psychometric curves and used them to simulate changing magnitudes of alteration for the SoR rating task based on their predicted detection probability: 25%, 50% (the Just Noticeable Difference or JND), 75%, and 95% (Fig. 1C). Unaltered trials were defined as having ‘0’ magnitude, which served as our baseline condition.

### SoR rating task

In the SoR rating task (Fig. 1D), participants were exposed to 360 trials of pseudo-randomly ordered conditions, 72 of which were baseline trials that did not include a virtual hallucination (magnitude 0). Each trial included a single appearance of the virtual environment for 5.2 seconds, during which participants performed the secondary task as instructed and observed the state of the environment. At the end of each trial the virtual environment disappeared followed by a gray question screen during which the participants were instructed to rate their experience in the virtual room (compared to the baseline condition) on a scale from “normal” to “strange” (100-0 respectively) using the VR controller. To control for potential drifts in ratings as a result of the participants losing touch with the baseline condition, we placed 5 baseline trials every 40 virtual hallucination trials. Participants were informed of these trials in advance by a message screen and were not instructed to rate them.

### Instructions & training for SoR rating task

Participants were instructed on the task, then entered the environment to refamiliarize themselves with the unaltered virtual environment. Training consisted first of the participants performing 9 trials using only 4^th^ magnitude stimuli for each condition. The rest of training consisted of an additional 18 randomly ordered trials simulating the actual task. Following training, participants immediately commenced the SoR rating task. An intermittent break was offered half way through to avoid fatigue. To avoid VR motion sickness, participants were instructed to report any discomfort and could stop the experiment at any stage.

### Clinical measures

After the experiment, participants completed three self-report questionnaires measuring symptoms related to SoR and anomalous perceptions: the Cardiff Anomalous Perceptions Scale (CAPS) (65), the Prodromal Questionnaire Brief Version (PQ-B) (66), and the Inventory of Psychotic-Like Anomalous Self-Experiences (IPASE) (67).

### Data exclusion criteria

Behavioral data were excluded post-data collection and before analysis. Pre-registered exclusion criteria included participants who did not understand the task or did not comply with the experimenter’s instructions prior to or during the experiment or if the participant was unable to finish the experiment. Trials with missing answers were excluded from analysis. Wrong answers were undefined in our data, we accepted all subjective judgments as reasonably valid answers.

However, we did require a minimal indication that participants’ SoR was affected by the task, so participants who showed no numerical decrease in average ratings between the baseline condition and the 4^th^ magnitude of alteration across all conditions would be excluded from further analyses.

Analysis of kinematics excluded trials with no movement or multiple movements as well as trials with early or late movements, and trials in which the hand was not returned to its initial position. Subsequently, participants with over 10% excluded trials were also excluded from analysis. A participant’s ECG data were excluded if more than 100 trials were considered outliers (mean ± 2 SD) or if the recording was faulty. Trials were excluded from pupil size analysis if the participant blinked during the first 100 ms of the trial. A participant’s pupil data was fully excluded if they had less than 8 trials per condition after removing unusable trials or if more than 20% of their data points needed interpolation.

### Statistical analysis

All statistical analyses were performed in R 4.4.1 and Python 3.9. Non-parametric tests are reported in cases where normality assumptions were violated. For mixed effects models, we used the lme4 (68) and robustlmm (69) libraries with effects reported using Welch’s ANOVA or Wald’s test. For Experiment 2 all statistical tests were one-sided based on our pre-registered hypotheses unless otherwise specified.

### Behavioral measures

The effects of virtual hallucination magnitude and domain on SoR ratings were analyzed using a linear mixed effects model. Domain-specific response patterns were quantified using slopes derived from the random effects of the linear mixed effects model. Additionally, we examined correlations between clinical scores and SoR metrics (average ratings and slopes per domain) using Spearman correlation, with outliers removed using the standard deviation method (mean ± 2 SD).

### Physiological measures

Based on the evidence that virtual hallucinations affected phenomenological judgments of reality, we examined whether they also influenced physiological measures. Specifically, we investigated if being exposed to altered reality modifies kinematic, pupillometric, or cardiac responses. To facilitate this analysis across all three signals, we employed a subgroup approach. We categorized trials by objective presence of virtual hallucinations (*altered* vs. *unaltered*), by objective strength (*unaltered* [magnitude 0], *weak* [1^st^ & 2^nd^ magnitudes], and *strong* [3^rd^ & 4^th^ magnitudes]), and by subjective experience (*normal & strange*, based on the 1^st^ and 4^th^ quartiles, respectively, of the min-max normalized ratings per participant. This strategy enabled us to detect potential differences in physiological patterns associated with varying degrees and perceptions of reality alterations. We conducted two types of analyses for these measures: hypothesis driven statistics, reported first, demonstrate the strength and reliability of physiological responses, and complementary machine learning classifications reaffirm the robustness of our pre-registered analyses, and demonstrate the potential to classify and predict SoR using complex signal combinations (see Classification of physiological signals).

### Kinematics

The kinematic data of 60 participants were analyzed (*n* = 30 from Experiment 1 and *n* = 30 from Experiment 2). Three participants were excluded for having insufficient data (over 10% of trials excluded) based on our pre-registered criteria (Supplementary Materials “Data exclusion criteria”). Velocity measures were computed from the positional data. The hand movements were segmented into two phases: reaching and retracting. The reaching phase began with movement initiation toward the butterfly and ended when the velocity on the Y-axis reached 0 m/s. The retraction phase was determined using the same criteria.

### Pupillometry

The pupil size data for 59 participants were analyzed (*n* = 30 from Experiment 1 and *n* = 29 from Experiment 2). Four participants were excluded for insufficient data (less than 8 trials per condition) based on our pre-registered exclusion criteria. We chose our main time window of interest between the 4^th^ and 5^th^ seconds after trial initiation (70–72). This allowed us to capture the delayed peak in pupil responses while ensuring pupil stabilization following luminance transitions between the virtual environment and the gray question screen.

### Heart rate

The heart rate data from 47 participants were analyzed (*n* = 24 from Experiment 1 and *n* = 23 from Experiment 2). We excluded 16 participants from analysis due to insufficient data (over 100 excluded trials) based on our pre-registered criteria. QRS complexes were detected and R-R intervals were used to calculate heart rate. Heart rate trajectory was down-sampled and averaged into 12 epochs of 430 ms. Heart rate change trajectory was computed as the difference of each epoch from the average of heart rate recorded in the 500 ms pre-trial period.

In the exploratory analysis of the cardiac data from Experiment 1, we observed distinct patterns in the averaged heart rate trajectories for trials with different subjective experiences (trials defined as *normal* or *strange* based on the 1^st^ and 4^th^ quartiles of min-max normalized ratings) and for trials from different domains. These observations led us to focus our analysis on time-locked heart rate responses to trial content using the averaged heart rate trajectory as our primary measure. To account for both the within-subject repeated measures nature of our data and the between-subject variability in heart rate responses, we analyzed the data using mixed effects models.

### Classification of physiological signals

To assess physiological markers of altered reality, we applied a machine learning logistic regression method using Scikit-learn (73) and corrected imbalances in the target variable data using oversampling from Imblearn (74) to predict *altered* or *unaltered* trials for each subject in the exploration experiment. Classification performance was evaluated using area under the curve (AUC) and F1 scores averaged across iterations. We generated 1000 models for each subject, with random train-test partitions. The models utilized an L2 penalty, C=10, and a max iteration of 1000. We ran the model on each signal separately, and on each signal combination. For cross-experiment validation, we created a composite model by averaging coefficients and intercepts across individual models from Experiment 1, which was then tested on data from Experiment 2. Data visualization and statistical reporting follow standard conventions, with error bars representing 95% confidence intervals.

### Computational model

As introduced earlier, SoR can be understood as a probabilistic inference process, where sensory inputs are evaluated against prior knowledge of the world. As a result, alterations in aspects of reality which we normally experience as inflexible such as the force of gravity should have a stronger impact on SoR (i.e., narrow likelihood distribution) than those we frequently experience in varied contexts such as color perception (i.e., wide likelihood distribution). To formulate this process, we modified a Bayesian causal inference model adapted from Chancel, Ehrsson & Ma (2022), commonly used in multi-sensory perceptual decision tasks (13,75,22). The model contains three key variables: the category *C*, the stimulus *x*_*trial*,_ and σ the sensory noise assumed by the observer. The category of the stimulus has a value of *C* = Real if *x*_*trial*_ was real or *C* = Unreal if *x*_*trial*_ was unreal. The prior probability of an event being judged as real before any sensory stimulation is *p*(*C* = Real) = *p*_*real*_.

On each trial the model observer receives a stimulus *x*_*trial*_ and infers its category *C* by computing the posterior probabilities *p*(*C* = Real | *x*_*trial*_) and *p*(*C* = Unreal | *x*_*trial*_). The comparison of the two posteriors is expressed as the log posterior ratio *d*. When there is equal evidence for both posteriors *d* = 0. In turn, the decision rule states the observer would infer *x*_*trial*_ is real when *d* > 0.

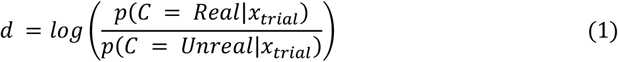

Importantly, while the posterior for reality, *p*(*C* = Real | *x*_*trial*_), is shaped by priors and likelihoods, the posterior for unreality in our model *p*(*C* = Unreal | *x*_*trial*_), represents a more diffuse, non-specific belief that any stimulus could be unreal, especially those at extreme values (Fig. 4A). This means that even familiar stimuli, such as a blue sky at noon, could theoretically be explained as non-veridical (e.g., hallucination, simulation), though in most cases, the posterior probability for the real explanation far exceeds that of the unreal one. The log-posterior ratio can be rewritten as a combination of the prior and likelihood ratios:

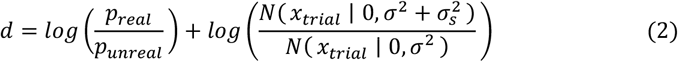

Assuming the observer infers *x*_*trial*_ is real when *d* > 0, we can rewrite this case of *d* in terms of *x*_*trial*_ like so:

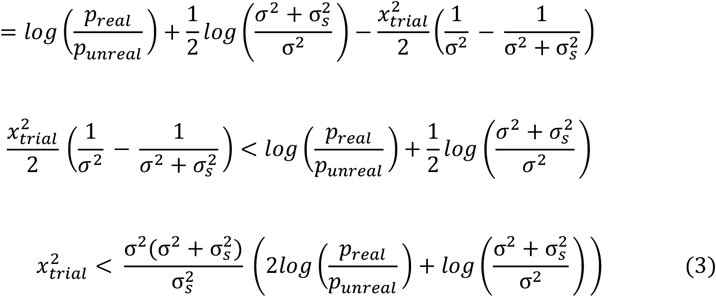

The right side of Eq. 3 is then defined as K. In the case where K > 0, the decision rule *d* > 0 is equivalent to → |*x*_*trial*_| < *k*, (Fig. 4A).

We fitted this model to participants’ SoR ratings across different domains and magnitudes. Since all participants experienced stimuli at five magnitudes corresponding to their individual psychophysical performances, the model accounted for these variations. In this model the response depends on the prior probability that *x*_*trial*_ is real (*p*_*real*_) and the sensory noise assumed by the observer in each trial (σ) (Fig. 4B). The model has two more free parameters: λ the lapse rate accounting for random guesses or unintended responses and we allowed an individual-level bias in using the SoR scale, by including a bias parameter that accounts for differences in average ratings between participants. The resulting model formula:

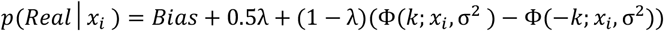

We then fitted the model to best capture the participants’ ratings, independently for each participant and each domain, using a root mean squared distance (RMSD) as a measure of fitting error:

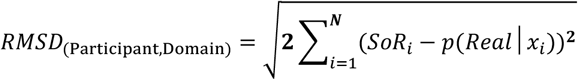

The *optim* function was used to find the set of parameters {λ, *Bias, p*_*real*_, σ} that minimized the RMSD error function for each participant in each domain. The output of the model is the probability of the observer deciding that *x*_*trial*_ is real.

### Model Fitting

For model fitting, we transformed the magnitude of stimuli from a linear scale of detection thresholds [0, 0.25, 0.5, 0.75, 0.95] into a normalized scale of reality-signal strength levels [0.05, 0.45, 0.55, 0.65, 0.95]. The units of the signal levels are arbitrary, reflecting the non-linear relations between signal strength and accuracy. The baseline magnitude for the model was set to a non-zero value adding flexibility to the estimates and also reflecting an inherent uncertainty on all trials attributed to task demands.

## Funding

This research project has received funding from an ERC starting grant (UNREAL-949010) to Roy Salomon.

## Author contributions

Conceptualization: RS

Formal analysis: GD, PBT, OH, AB

Funding acquisition: RS

Investigation: GD, PBT, OH, AB,

NG Software: YZ

Methodology: RS, AZ, UH, GD, PBT

Visualizations: GD, PBT, OH, AB

Writing – original draft: RS, GD

## Competing interests

The authors declare no conflict of interest.

## Data and materials availability

All data needed to evaluate the conclusions in the paper are present in the paper or the Supplementary Materials. Data, analysis code, and materials will be made publicly available on GitHub at: https://github.com/SLabhaifa upon publication.

## Notes

### Competing Interest Statement

The authors have declared no competing interest.

### Summary of Updates

Changed the text in the title page

