## Supplementary material for "Unreal? A Behavioral, Physiological, and Computational Model of the Sense of Reality"

Gadi Drori *et al.*

**This PDF file includes:**

Supplementary Text
Figs. S1 to S4
Tables S1 to S2
References (76 to 82)

### Supplementary Text

Power Analysis

To calculate the required sample size for the Experiment 2 experiment we used Kendall's $W$ effect size for a Friedman test performed on the data from Experiment 1. We based our calculations on an evaluation of differences in reality ratings between magnitudes of alteration (across conditions). Kendall's $W$ interpretation guidelines are 0.1 - 0.3 (small effect), 0.3 - 0.5 (moderate effect), and > 0.5 (large effect). A large effect size was detected for the difference between ratings across condition magnitudes ($W = 0.95$). We then used the G*Power software v3.1.9.4 (76) to calculate the required sample size for our Experiment 2 study with the following parameters: effect size $W = 0.95$, alpha = 0.0001, power = 0.95. The calculated sample size based on these parameters was $N = 32$.

Hardware and software specifications

All experiments were performed on an Intel core i9 processor and 32 GB of RAM computer with an NVIDIA RTX 3070 graphics card, running our in-house software UnReal-VR, built with Unity 2018.3.2. During the experiments participants wore the HTC Vive-pro Eye head-mounted display with a resolution of 2800 × 1600 pixels, a refresh rate of 90 Hz, eye-tracking capabilities and retractable headphones. Participants responded using the HTC Vive touch-sensitive hand controller. Motion tracking was performed using the HTC Vive (2.0) base stations. All VR hardware was manufactured by Valve Corp., Washington, DC, USA.

VR eye-tracking

To ensure the quality of pupil data each time a participant donned the VR helmet or made significant adjustments to it, they underwent a built-in three step eye tracking calibration procedure. To minimize external interference with pupil size due to auditory interferences, participants listened to pink noise using the HMD headphones. It is well-documented that auditory stimuli can influence pupil size, necessitating the use of consistent background noise to ensure accurate measurements (77).

ECG Harness

Electrocardiography (ECG) signals were acquired using the Biopac Bioharness device (BIOPAC Systems, Inc., Goleta, California) with two electrodes connected via Bluetooth at a rate of 250 Hz.

Data streams were synchronized through a LAN network setup using Lab Streaming Layer (LSL) (78).

Virtual Environment

We adapted the ArchVisPro Interior Vol.4 virtual environment from the Unity Asset Store with real depth and width processing, chosen due to its highly immersive and realistic appearance, representing our normative space modeled as a house with furniture.

Setup and calibration procedures

Participants were seated in the experiment room in front of a table, holding the VR controller in their right hand. Participants were instructed to keep their controller hand at the center of a black X mark on top of the table with a square outline and raised texture added for increased tactile feel. This mark served as the hand's start and finish position each trial. In VR, participants were placed at the center of the virtual house where they could observe and interact with the environment from a seated position. We matched several virtual objects with real objects to create the illusion of a physical correspondence between them, thus increasing the immersiveness of the simulation. Participants sat in front of a virtual table matching in height, location and the hand position marker to the real table in front of them. In addition, the participant's height while seated upright was measured using the VR headset before starting the experiment to ensure the distance between their head and the real table corresponds to the distance between their virtual first-person perspective (1PP) and the height of the virtual table. The hand holding the VR controller was represented using the RRFreelance Animated Hands with Gloves 2019.3 HDRP virtual asset, including a representation of the HTC Vive controller with corresponding finger movements. At the opposite edge, on top of the table, sat a glass terrarium housing a hidden butterfly and above the terrarium on the ceiling hung a planter with flowers.

Detailed secondary task

The default scene for all experimental tasks consisted of the hanging plant pot falling 1.5 seconds after the start of each trial (with a jitter of ± 50 ms). Each time the pot falls it hits the top of the terrarium triggering the hidden butterfly to fly upwards and reveal itself. Participants were instructed to perform a reaching movement with their right hand towards the location of the butterfly as soon as it was visible, then wait for the butterfly to disappear again, and return their hand to the X mark to complete the trial.

The aim of this experimental design was to provide opportunity on each trial for participants to detect changes in the environment across our three domains of experience. The dynamics of the scene and the instruction to observe specific events and respond to them requires participants to sample the visual appearance and physics of the environment and their own first-person perspective. After each trial, the virtual room was replaced by a gray question screen, with a sliding scale ruler ranging from "normal" to "strange" with which participants used to report their answers before proceeding to the next trial.

Detailed staircase procedure

Participants went through 9, two-alternative-forced-choice (2AFC) staircase procedures in succession for all virtual hallucination conditions (the order of the staircases was randomized between participants). In the 2AFC procedure, participants were successively immersed in the virtual environment in two phases (for 5.2 seconds each), with a black screen displayed (0.5 sec) between them. In each phase, the environment was identical except that one phase included one of the nine virtual hallucinations. After viewing the two phases, participants were presented with a gray question screen with a sliding bar where they were required to answer using the VR controller touch pad whether the two phases were "different" (press left) or "the same" (press right) indicating if they were able to detect the alteration.

We employed a 1-up\1-down descending staircase to sample psychophysical sensitivity and derive detection thresholds for each virtual hallucination. Since staircases were randomly ordered, each new staircase was presaged with a "new condition" message screen and the first three trials of the new staircase were intentionally set to extreme values and defined as throw-away trials, to make sure that the participants were fully aware of the virtual hallucinations being presented to them. During throw-away trials, the experimenters were allowed to quickly halt the procedure and provide some direction in case the participant did not report noticing the virtual hallucinations. The magnitude of the parameter controlling the virtual hallucinations was decreased exponentially by a factor if the participant reported the last two phases were "different" and was increased by the same factor if they were reported as "the same". The stopping rule for the procedure was defined after 5 reversal points or 25 trials in either direction. In case the data collected for a given condition was insufficient to model psychophysical thresholds we repeated the procedure for that condition, but this time starting the staircase procedure one step higher than before (including the throw-away trials) to facilitate initial detection. To avoid exhausting the participants with prolonged VR sessions, we excluded subjects that required more than 18 staircase procedures total (regardless of which conditions were repeated). The data collected in session 1 enabled the calculation of individually fitted parameter values based on the participant's sensitivity to each virtual hallucination condition in units based on their Just Noticeable Differences (JNDs). This allowed us to match the perceptual saliency of the stimuli within and between participants for the SoR rating task.

Detailed curve estimation procedure

We fit psychometric curves to the response data using the Psignifit toolbox in MATLAB, implementing a logistic function of the form:

$$p(\mathrm{x}) = \gamma + (1 - \gamma - \lambda)/(1 + \exp(-2\log(1/\alpha - 1)/\beta(x - m)))$$

Where $x$ is the stimulus magnitude, $m$ is the threshold, and β is the width parameter. The response probability $p$(different) for a given stimulus magnitude was modeled with four parameters: threshold ($m$), width (β), guess rate (γ), and lapse rate (λ). We performed Bayesian inference using Maximum a-posteriori (MAP) estimation of the psychometric curve parameters. Model fits were evaluated using McFadden's pseudo-$R^2$.

We implemented additional conservative criteria for fit validation: (1) we required at least one sampled value to fall on the slope of the psychometric function (defined within the 15%-85% response range), and (2) we established a stringent minimum threshold of $R^2 \geq 0.3$ for fit acceptance otherwise, the staircase for that condition was repeated until the fit was sufficiently good. These criteria, while potentially excluding some valid fits, were deliberately conservative to ensure reliable conclusions despite the limited sampling necessitated by experimental time constraints. In Experiment 2 we added a requirement to reject $R^2 > 0.99$ which were associated with an insufficient variance in sampling and responses.

Detection thresholds for each of the nine conditions were estimated from the fitted data at 4 points along the psychometric curves: 25%, 50% (the JND), 75%, and 95%. The threshold values served as the orders of magnitude for each virtual hallucination in the SoR rating task (magnitudes 1 - 4 respectively). We constrained threshold estimates to within the range of sampled stimulus magnitudes. For values exceeding this range, the highest threshold ($4^{th}$ magnitude) was capped at the maximum sampled value, and the $3^{rd}$ magnitude, if out of range, was set to the mean of the $2^{nd}$ and $4^{th}$ magnitudes. The two lowest threshold magnitudes remained unadjusted.

Kinematic data preprocessing

Hand movement data underwent preprocessing, including cleaning and interpolation. Velocity was computed using the Euclidean norm of positional derivatives along the x, y, and z axes. Time intervals between data points were corrected to a consistent 11 ms via cubic interpolation using SciPy (79). To maintain a fixed trial length of 5.2 seconds, each trial was adjusted to contain exactly 473 data points. This number was derived from the sampling rate of the VR equipment (5200 ms / 11 ms ≈ 472.72, rounded up to 473). If necessary, trial lengths were corrected by subtraction from or addition to the end of the trial.

Identifying kinematic points of interest

Points of interest in the arm movement were derived to facilitate further analysis. These points of interest served as markers for movement segments that were consistently present across trials, enabling the identification and exclusion of specific trials in which arm movements were not successfully completed. One crucial landmark is the movement initiation threshold, which indicates the beginning of a movement. In this study, the threshold velocity defining movement initiation was 0.001 meters per second on the Y axis. The Y axis was chosen due to the significant variation of the data on this axis observed across trials. The arm movement was segmented into two phases: reaching and retracting. The start of the movement represented the initiation of the reaching phase toward the butterfly, which ends when the velocity on the Y axis reaches 0, indicating the completion of the reaching motion. The retraction phase was determined using the same criteria.

ECG data preprocessing

Preprocessing of the ECG signals was performed using Neurokit2 (80). The preprocessing pipeline included a 0.5 Hz high-pass Butterworth filter (order = 5) to remove low-frequency noise and subsequent power line filtering at 50 Hz. QRS complexes were detected by evaluating the steepness of the ECG signal's absolute gradient, with R-peaks identified as local maxima. The resulting R-R intervals were used to calculate heart rate. To facilitate further analysis, heart rate was down-sampled and averaged into 12 epochs of 430 ms duration per trial (5200 ms / 12 = 433.33).

Pupil preprocessing

Pupil size data were recorded at a frequency of 90 Hz directly from the VR headset and extracted using the SRanipal software development kit v1.3.3 provided by HTC (81). We defined the relevant time period for our analysis between the 4th and 5th seconds from the start of each trial and concatenated the time series by trial. This time window was chosen to ensure sufficient trial time for all alterations to be detected and for pupil size to stabilize after luminance transitions originating at the start of the trial (from the question screen to the virtual environment). The mean time-series for each magnitude can be seen in Figure S4B-C. Blinks were detected and marked automatically by the HTC Vive eye-tracker. To further guarantee that blinks did not confound pupil size in these segments a 55 ms time window surrounding each blink was removed from the data (82). Trials in which we had to remove more than 20% of the data were excluded to allow a more reliable interpolation of the remaining data. The signal was smoothed (Hamming = 10), interpolated and sampled every 11 ms using SciPy, resulting in a continuous time series. To control for pre-trial changes in pupil size the mean of the first 100 ms for each trial was subtracted from the entire trial. The Perception domain included alterations that could potentially influence pupillometry in unknown ways (saturation, ripple). To ensure the accuracy of our analysis, we preregistered and excluded all trials associated with this domain from the pupil analysis.

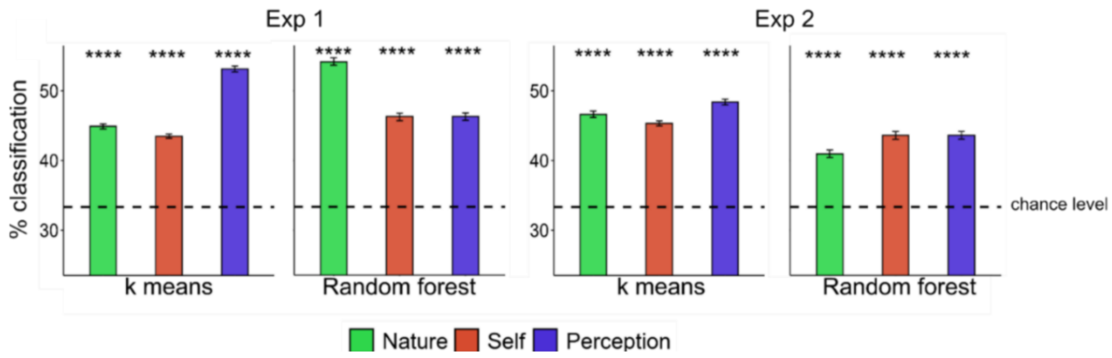


**Fig. S1. Data-driven classification of ratings by domain.** We performed k-means and random forest classifications (n = 1000 each) to validate the data from different domains (Nature, Self and Perception) indeed represent coherent behavioral foci (i.e., virtual hallucinations within a domain are more similar than across domains). We successfully classified SoR ratings to virtual hallucination domains above chance level in both experiments (all $p$ values < 0.001, corrected). Bars and error bars represent mean ± bootstrapped 95% CI, dashed line represents chance level classification (33.3%). The classification proportions for each domain were compared to the chance level with two-sided Wilcoxon signed rank tests. K-means yielded more consistent effect sizes ($r = 0.87$) than random forest ($0.69 \leq r \leq 0.86$), with comparable performance between experiments.

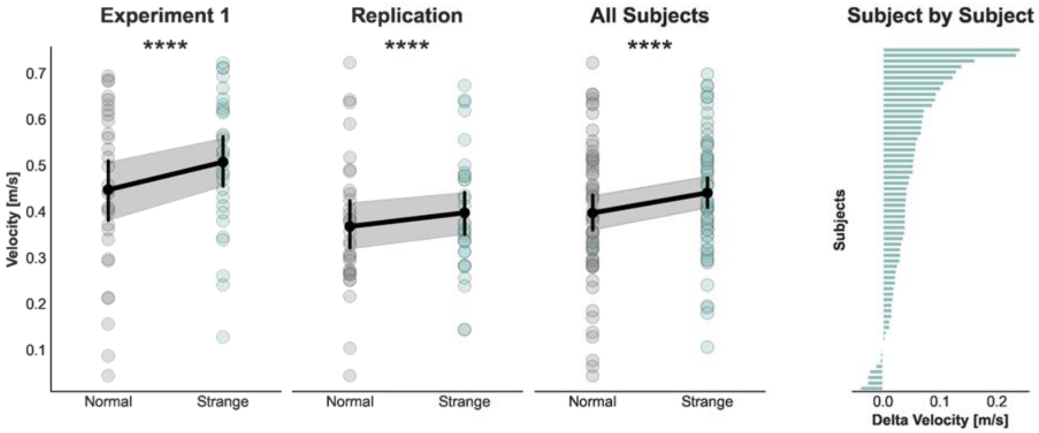


**Fig. S2. Hand velocity increases in subjectively strange trials.** Exploratory analysis revealed hand velocity during the reaching phase of strange trials was 17.8% higher (0.48 ± 0.16) than in normal trials (0.45 ± 0.17) in Experiment 1 ($W_{29}$ = 33, $p < 0.0001$). Hand velocity was 13.82% faster in strange trials (0.39 ± 0.14) than in normal trials (0.37 ± 0.15) in Experiment 2 ($t_{29}$ = -5.11, $p < 0.0001$).

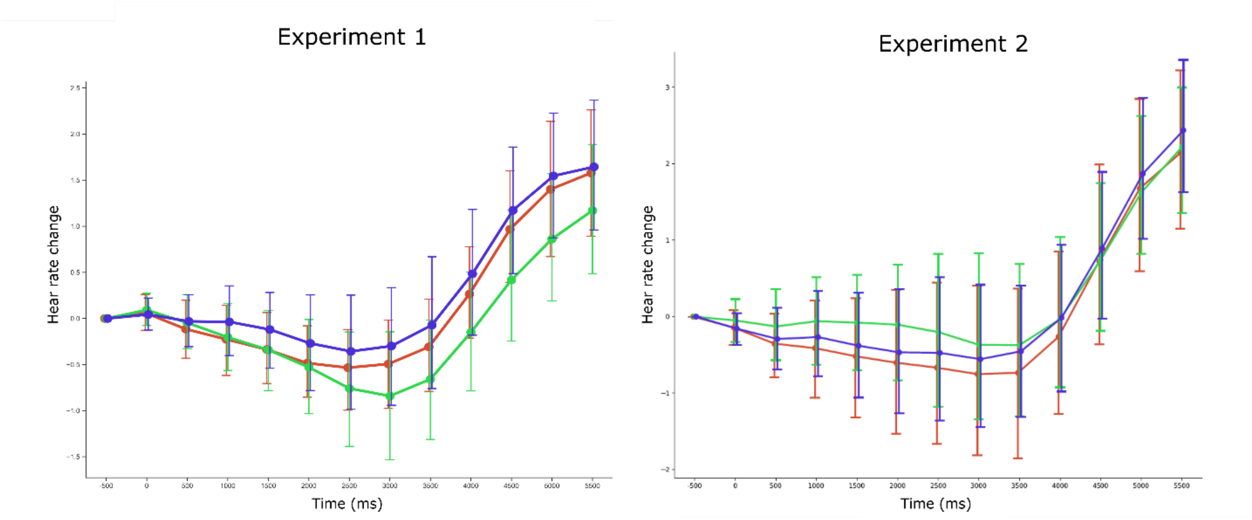


**Fig. S3. Cardiac deceleration during strongly altered trials.** Heart rate deceleration differed between domains in strong trials in Experiment 1 ($\chi^2_2$ = 14.63, $p < 0.001$, $d$ = 0.78), but not in Experiment 2 ($\chi^2_2$ = 5.81, $p$ = 0.054, $d$ = 0.49) [H2.e ×].

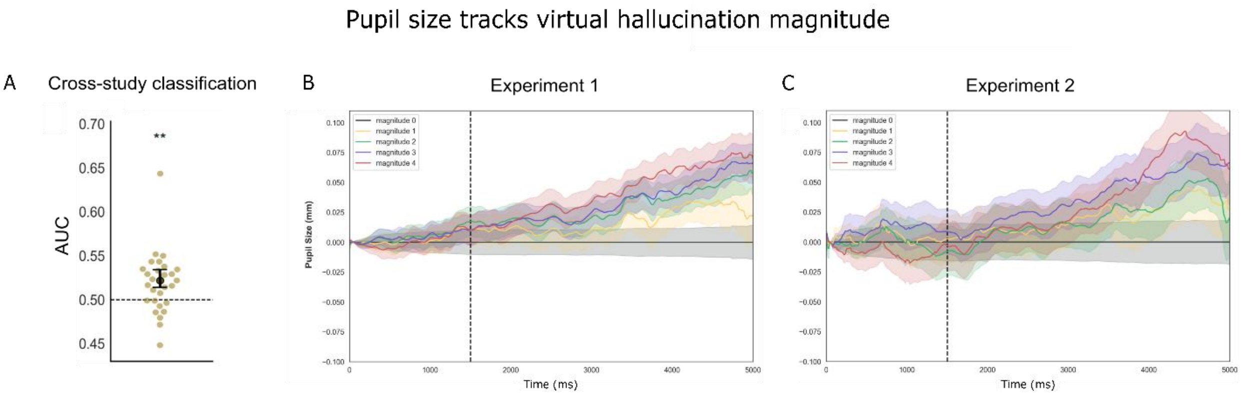


**Fig. S4. Pupil dilation tracks alteration magnitude. (A)** A classification model trained on data from Experiment 1 successfully predicted virtual hallucination presence above chance level using pupil size data from Experiment 2 (AUC = 0.53 ± 0.03, $t_{28} = 4.99$, $p < 0.001$, $d = 1$). **(B-C)** Average pupil size increased with virtual hallucination magnitude in both experiments.

**Random Effects Comparison**

| Model | npar | AIC | BIC | logLik | Deviance | Chisq | Df | p_value | Delta_AIC | Delta_BIC |
|---|---|---|---|---|---|---|---|---|---|---|
| Intercept by subject | 17 | 3737.7 | 3808.1 | -1851.8 | 3703.7 | NA | NA | NA | NA | NA |
| Full model | 22 | 3637.5 | 3728.7 | -1796.8 | 3593.5 | 110.12 | 5 | < 2.2e-16 | 100.2 | 79.4 |
| Intercept by domain | 17 | 3909.8 | 3980.2 | -1937.9 | 3875.8 | NA | NA | NA | NA | NA |
| Full model | 22 | 3637.5 | 3728.7 | -1796.8 | 3593.5 | 282.24 | 5 | < 2.2e-16 | 272.3 | 251.5 |

**Table S1. Comparison of random effects.** We analyzed the effects of virtual hallucination magnitude and domain on SoR ratings of Experiment 1 using a two-step model selection approach. First, to determine the optimal random effect structure, we compared three mixed-effects models using likelihood ratio tests. First, we compared a model with only random intercepts for participants (intercept only model) to a model with both random intercepts and slopes for domain by participant (full model). The model including random slopes (full model) provided a significantly better fit, as indicated by a substantial reduction in AIC (ΔAIC = 100.2, ΔBIC = 79.4) and a highly significant likelihood ratio test ($\chi^2_5 = 110.12$, $p < 0.0001$). This suggests that allowing individual differences in domain effects per participant improves explanatory power. Next, we compared the full model to a model with only domain as a random intercept (intercept by domain). The full model again showed a substantially better fit (ΔAIC = 272.3, ΔBIC = 251.5, with a highly significant likelihood ratio test ($\chi^2_5 = 282.24$, $p < 2.2e\text{-}16$). This confirms that individual variability in domain effects across participants is a crucial factor to account for. Therefore, we selected the model including both random intercepts and slopes for domain by participant (SoR rating ~ Magnitude * Domain + (Domain | Subject).

Fixed Effects Comparison

| Model | Intercept | Domain | Level | Interaction | df | logLik | AICc | Delta_AICc | Weight |
|---|---|---|---|---|---|---|---|---|---|
| Full Model (Interaction) | 87.35 | + | + | + | 16 | -1937.892 | 3909.0 | 0.00 | 0.688 |
| Without Interaction | 81.48 | + | + | | 8 | -1947.134 | 3910.6 | 1.58 | 0.312 |
| Model 3 | 87.17 | | + | | 6 | -1965.710 | 3943.6 | 34.60 | 0.000 |
| Model 4 | 61.41 | + | | | 4 | -2064.965 | 4138.0 | 229.02 | 0.000 |
| Model 5 | 67.10 | | | | 2 | -2076.332 | 4156.7 | 247.69 | 0.000 |

**Table S2. Comparison of fixed-effects.** Next, we used an automated model selection method *dredge* (MuMIn) to compare all possible fixed effects combinations. This revealed that the full model including domain, magnitude, and their interaction provided the best fit (AICc = 3909, weight = 0.688), followed by a model without the interaction (ΔAICc = 1.58, weight = 0.312).
